# Polarity remodeling of ventricular cardiomyocyte precursors during heart morphogenesis is required for ventricular chamber formation

**DOI:** 10.64898/2026.05.20.726717

**Authors:** Zihang Wei, Jingzhen Chen, Chuyao Peng, Xiaotong Wu, Anming Meng

## Abstract

Vertebrate heart development requires precise regulation of morphogenetic movements and dysregulation causes congenital heart defects. However, cell polarity remodeling during early heart tube formation remains underexplored. Here, we show that zebrafish ventricular cardiomyocyte precursors (vCPCs) undergo striking polarity reorientation as they transition from a single-layered epithelium to a transient double-layered configuration by involution. During this process, vCPCs remove inner adherens components and re-establish polarity toward the ECM-attached outer surface, enabling proper directional extension to form the ventricle. We identify the junctional scaffold protein Afadin a (Afdna) as a critical regulator. Afdna localizes to the outer and lateral membrane to restrict Rap1 activity, facilitating Podocalyxin translocation and adhesion disassembly at the inner interface between vCPC double layers. *afdna* mutants exhibit a multilayered ventricle with impaired blood flow, phenocopying human congenital ventricular obliteration. Our findings uncover a polarity-based mechanism ensuring ventricular chamber formation and provide insight into ventricular obliteration.

## Introduction

Organ morphogenesis is a highly orchestrated process driven by the dynamic behaviors of cells. Understanding these cellular dynamics is crucial, as their disruption often underlies congenital disorders^1^. This is particularly true in the heart, where congenital heart diseases (CHD) such as single ventricle defects^2–4^, hypoplastic left heart syndrome^5^, hypertrophic cardiomyopathy^6,7^, and dilated cardiomyopathy^8,9^ are usually linked to structural abnormalities of developmental origin. However, cellular and molecular mechanisms underlying CHD during heart morphogenesis are underexplored. Although mammalian models, particularly mice, have been instrumental in identifying genetic regulators of cardiogenesis^10,11^, a fundamental understanding of the precise cellular rearrangements that build the heart tube has been hindered by technical challenges of live imaging in utero.

The zebrafish, with its optical clarity and high genetic conservation to mammals, provides a powerful model to visualize and dissect these dynamic morphogenetic events^12^. The conservation of key developmental genes and pathways makes zebrafish a suitable system to model the cellular and molecular etiology underlying certain CHDs^13,14–16^. Early heart development in zebrafish embryos proceeds through a series of conserved stages: specification, migration, involution, and tube extension^17,18^. Cardiomyocyte progenitor cells (CPCs), including ventricular CPCs (vCPCs) and atrial CPCs (aCPCs), are specified from the anterior lateral plate mesoderm (ALPM), and start to migrate towards the midline as a single-layer epithelial sheet at the 14-somite stage. Upon reaching the midline around the 18-somite stage, several vCPCs adjacent to the midline involute to form a double-layered cardiac cone^13^. This transient double-layered structure subsequently tilts and extends along the anterior-posterior axis, guided by left-right asymmetric cues such as Lefty^19,20^, ultimately forming the single-layered linear heart tube.

Many critical morphogenetic events require precise regulation of cell polarity, such as neural tube formation^21^, lumen formation^22^, and epithelial formation^23^. Previous studies have established that during heart development in zebrafish embryos, CPCs within the double-layer cone exhibit polarized characteristics, and disruptions in this polarity are linked to severe cardiac developmental defects^24–27^. Key junctional and adhesion molecules, including zonula occludens protein ZO-1^28^, atypical protein kinase C (aPKC)^29^, and β-catenin^28^, are asymmetrically localized in the polarized cells. Furthermore, genetic studies in mice confirm that disrupting cardiomyocyte polarity leads to developmental defects^30–32^, underscoring an evolutionarily conserved role for polarity in heart formation. Despite these advances, the regulatory mechanisms and functional significance of the dynamic polarity remodeling that must occur during the critical involution step in the zebrafish have remained largely unexplored.

In this study, we combine high-resolution imaging and genetic approaches in zebrafish embryos to define the mechanism of heart tube formation. We focus on the involution process, revealing how polarity reorientation is triggered by extracellular cues and executed by an intracellular signaling cascade. Our work identifies the junctional scaffold protein Afdna, an orthologue of mammalian AFDN, as a critical spatial regulator that orchestrates a GTPase-dependent pathway to ensure adhesion remodeling and successful tube extension, thereby elucidating a fundamental morphogenetic module in heart morphogenesis.

## Material and methods

### Zebrafish strains and maintenance

Wild-type zebrafish of the Tuebingen (Tu) strain were utilized for all experiments. Embryos were maintained in Holtfreter’s solution at 28.5°C and were collected at specific developmental stages. All zebrafish husbandry and experimental procedures were conducted in accordance with the guidelines set by the Animal Care and Use Committee at Tsinghua University.

### Transgenic and mutant zebrafish lines

In this study, we utilized several transgenic and mutant zebrafish lines. The transgenic lines *Tg(myl7:EGFP)* and *Tg(myl7:afdna-GFP)* were generated using the *Tol2* transposon system in WT and the transgene was introduced into *afdna^tsu-^*^5^ mutant background by breeding when necessary. The mutant line *afdna^tsu-^*^5^ was generated by CRISPR-Cas9 technology. Cas9 protein (NEB, M0386S) and sgRNA were injected into one-cell stage WT embryos, followed by screening. The sgRNA target sequence was 5’-GAGAATGTCGGGGAGCCGAGAGG-3’. sgRNA was transcribed by MEGA T7 kit (Thermo Fisher Scientific, AM1333), then purified by alcohol precipitation. To identify mutants, embryos from *afdna* heterozygote incrosses were genotyped by PCR after observation by microscopy. The primers used for PCR were 5’-CGATAAGACAAGCGAGCGACGG-3’ (forward primer), 5’- TCCGCCGCTCCTCCTCTCGG-3’ (reverse primer for WT allele) and 5’-TCCGCCGCTCCTCCTCTCCC-3’ (reverse primer for mutant allele).

### mRNA synthesis

The target coding sequence was cloned into pCS2 plasmid, and recombinant plasmid was identified by PCR. The linearized plasmid after restriction enzyme digestion was purified either through alcohol precipitation or with a DNA purification kit (Tiangen, DP214). For the synthesis of mRNA, different methods were employed based on the length of the transcript. Short mRNA transcripts (less than 4 kb) were generated using the mMessage mMachine SP6 Kit (Thermo Fisher Scientific, AM1340). Following transcription, the mRNA was purified using RNA purification magnetic beads (Vazyme, N412). For longer mRNA transcripts, the MEGAscript SP6 Kit (Thermo Fisher Scientific, AM1330) was used for transcription. The resulting mRNA was purified by lithium chloride precipitation. Subsequently, the purified naked mRNA was capped using the Capping System (Novoprotein, M082). The capped mRNA was then further purified using RNA purification magnetic beads (Vazyme, N412).

### Morpholino and mRNA injection

Morpholinos used in this study were purchased from Gene Tools LLC. The sequences and injection dose (per embryo) of the Morpholinos are as follows. *podxl*: 5’-GGTCATTTTCAGATTCTCCGCGTTC-3’ ^33^, TGGTGGCAGATTATTTCTTTTCACC-3’ ^34^, ACGCATTGTGCAGTGTGTCCGTTAA-3’ ^34^, CTTCTTGCGAATTGCTGCCATTTTG-3’ ^35^, 0.3 nM. Morpholinos were dissolved in nuclease-free water and diluted to the appropriate concentration for injection^36^. For mRNA injections, in vitro synthesized mRNA was also diluted in nuclease-free water. Morpholino or mRNAs were injected into the cytoplasm of 1-cell stage embryos. Injection doses of mRNAs were indicated in corresponding figure legends.

### Blood flow tracing

Embryos at 1 dpf or 2 dpf were anesthetized and placed on an injection plate. Rhodamine B-conjugated dextran (Thermo Fisher Scientific, D7139) was then injected into the common cardinal vein of the embryos. Following the injection, the embryos were transferred to Holtfreter’s solution containing MS-222 (Merck, E10521) for further anesthesia and immobilization. The embryos were subsequently embedded in low-melting agarose to facilitate positioning for confocal imaging. Blood flow was visualized and analyzed using confocal microscopy to assess circulation patterns and any potential abnormalities.

### Whole-mount immunofluorescence and confocal microscopy

Whole-mount immunofluorescence was performed following a standard protocol with modifications as needed. Briefly, embryos were dechorionated using protease treatment to remove the chorionic membrane. Embryos for immunofluorescence of ZO-1, Cdh2, Rab11, mCherry, α-catenin and β-catenin were fixed in 4% paraformaldehyde at room temperature for 4 h. Embryos for immunofluorescence of Afdna and fibronectin were fixed in freshly prepared methanol/DMSO (4:1) at 4°C overnight. Embryos fixed in methanol were first rehydrated through a graded series of methanol in PBSTx (75%, 50%, 25% and 0% methanol in PBSTx). Then, embryos fixed in either condition were permeabilized in 0.5% PBSTx (phosphate-buffered saline with 0.5% Triton X-100) for 30 min. The embryo was manually cut off at the midbrain-hindbrain border (roughly anterior to the cardiac mesoderm) using a scalpel (Extended Data Fig. 1). The trunk region of the dissected embryo was subjected to blocking and antibody staining according to standard procedures.

The primary antibodies used in this study include: mouse ZO-1 antibody (Invitrogen, 33-9100, 1:100 dilution); rabbit Cdh2 antibody (Abcam, ab211126, 1:100 dilution); rabbit Fibronectin antibody (Merck, F3648, 1:100 dilution); chicken GFP antibody (Abcam, ab13970, 1:400 dilution); rabbit mCherry antibody (Cell Signaling Technology, E5D8F, 1:200 dilution); rabbit Rab11a+b antibody (GeneTex, GTX127328, 1:200 dilution); rabbit Afdna antibody, custom-made by HuaBio using the target peptide PDYAPRKGRKPDNR. For secondary antibody and dye detection, the following reagents were used: goat anti-Chicken Alexa Fluor 488 (Abcam, Catalog No. ab150169, 1:200 dilution); multi-rAb CoraLite Plus 488 Goat Anti-Mouse (Proteintech, RGAM002, 1:200 dilution); multi-rAb CoraLite Plus 594 Goat Anti-Mouse (Proteintech, RGAM004, 1:200 dilution); multi-rAb CoraLite Plus 647 Goat Anti-Rabbit (Proteintech, RGAR005, 1:200 dilution); Phalloidin-iFluor 405 Reagent (Abcam, ab176752, 1:200 dilution); DAPI (Solarbio, C0060, 1:100 dilution).

The immunostained embryos were embedded in low-melting agarose, positioned appropriately, and imaged using the LSM980 Airyscan2 system (Extended Data Fig. 1).

### Whole-mount in situ hybridization and Fluorescence in situ hybridization (FISH)

Whole-mount in situ hybridization (WISH) for embryos was carried out following a standard protocol to detect specific gene expression patterns. Fluorescence in situ hybridization (FISH) and dual-color FISH combined with immunostaining were performed according to published methods^37^. The primers used for probe amplification in WISH and FISH included: *nkx2.5:* 5’-ATGGCAATGTTCTCTAGCCAAATG-3’ (forward primer) and 5’-TAATACGACTCACTATAGGGCCAAGCTCTGATGCCATGTAGTG-3’ (reverse primer); *myl7:* 5’-GTCCATGTAGGGGACGAACAG-3’ (forward primer) and 5’-TAATACGACTCACTATAGGGATTAACAGTCTGTAGGGGGCAG-3’ (reverse primer); *myh6:* 5’-AAGCCACTACCGCCTCTCTA-3’ (forward primer) and 5’-TAATACGACTCACTATAGGGAGTGCTTAAACTGCGAGCCT-3’ (reverse primer); *myh7:* 5’-ATTCTGAGGTGGCACAGTGG-3’ (forward primer) and 5’-TAATACGACTCACTATAGGGCTGCAGAAGGTTGTTGCGTC-3’ (reverse primer). Light-sheet imaging was performed using a MUVI-SPIM microscope.

### Western blotting

Zebrafish larvae at 5 dpf were collected into centrifuge tubes and washed three times with PBS. The larvae were then lysed using RIPA buffer with the aid of a mini grinder, and the lysate was incubated on ice for 30 min. The lysate was subsequently centrifuged and the supernatant was collected for Western blot analysis. Afdna antibody was described above, and mouse α-tubulin antibody (Proteintech, 11224-1-AP, 1:2000 dilution) was also used.

### Section and H**&**E staining

Embryo at 2 dpf was fixed in 4% PF overnight, Samples were embedded in paraffin wax, dehydrated in ethanol, and sectioned into 5-μm slices. Then the paraffin sections of different samples were stained with H&E and photographed under a microscope.

### Chemical treatments

Embryos at 8-somite stage (13 hpf) were manually dechorionated by needles, placed at 6-well plate with 3 ml filtered holtfreter’s solution containing GGTI-298 Trifluoroacetate (MCE, HY-15871), Narciclasine (MCE, HY-16563), or both, and incubated at 28.5℃ for 8 hours. Embryos then washed in fresh Holtfreter’s solution 3 times and raised at 28.5℃. Blood circulation was observed at 2 dpf, followed by genotyping of individuals.

### Statistical analysis

An average from multiple samples was expressed as mean ± SD (standard deviation). The significance of the difference between groups was analyzed by Student’s *t*-test (two-tailed). Significant levels were indicated in the corresponding context.

## Results

### Ventricular cardiomyocyte precursors (vCPCs) involute during their medial migration

Previous studies have demonstrated that the two lateral CPC domains, arranged as a monolayer sheet on each side of the midline, begin their medial migration at midsomitogenesis (around the 14-somite stage) and complete midline fusion to form the cardiac cone by approximately the 22-somite stage^19,28,29^. However, dynamic movements of CPCs are still poorly understood due to imaging technical and equipment limitations. We decided to re-analyze this process by light-sheet microscopy, which allows for high-speed, three-dimensional visualization of embryos with minimal photodamage^38^. Embryos at critical stages of the cardiac cone formation (16-somite stage to 22 hours postfertilization (hpf)) were subjected to dual-color fluorescence in situ hybridization (FISH) using probes for the pan-cardiomyocyte marker *cmlc2/myl7* and the ventricular-specific marker *vmhc/myh7* and then the developing cardiac cone was observed by light-sheet fluorescence microscopy (Fig. 1a and Extended Data Fig. 1a). The result showed that the leading cell of both left and right CPC sheets had involuted in most embryos at 18-somite stage (Fig. 1b, h), the number of involuted CPCs increased progressively as development proceeded (Fig. 1c-e, h). The involuted CPCs overlaid the non-involuted CPCs to form double layers with a maximum of four cells in the top layer before 23-somite stage. The involution movement was restricted to *myh7*-positive vCPCs since no *myh7*-negative atrial CPCs (aCPCs) were detected within the double-layered structure, which is consistent with previous reports^19,39^. Our observations clearly demonstrated concurrent medial migration and involution of both left and right vCPCs, which is in contrast to the previous model in which involution initiates only after the completion of the medial fusion and is restricted to one side^19,39^.

**Fig. 1.**
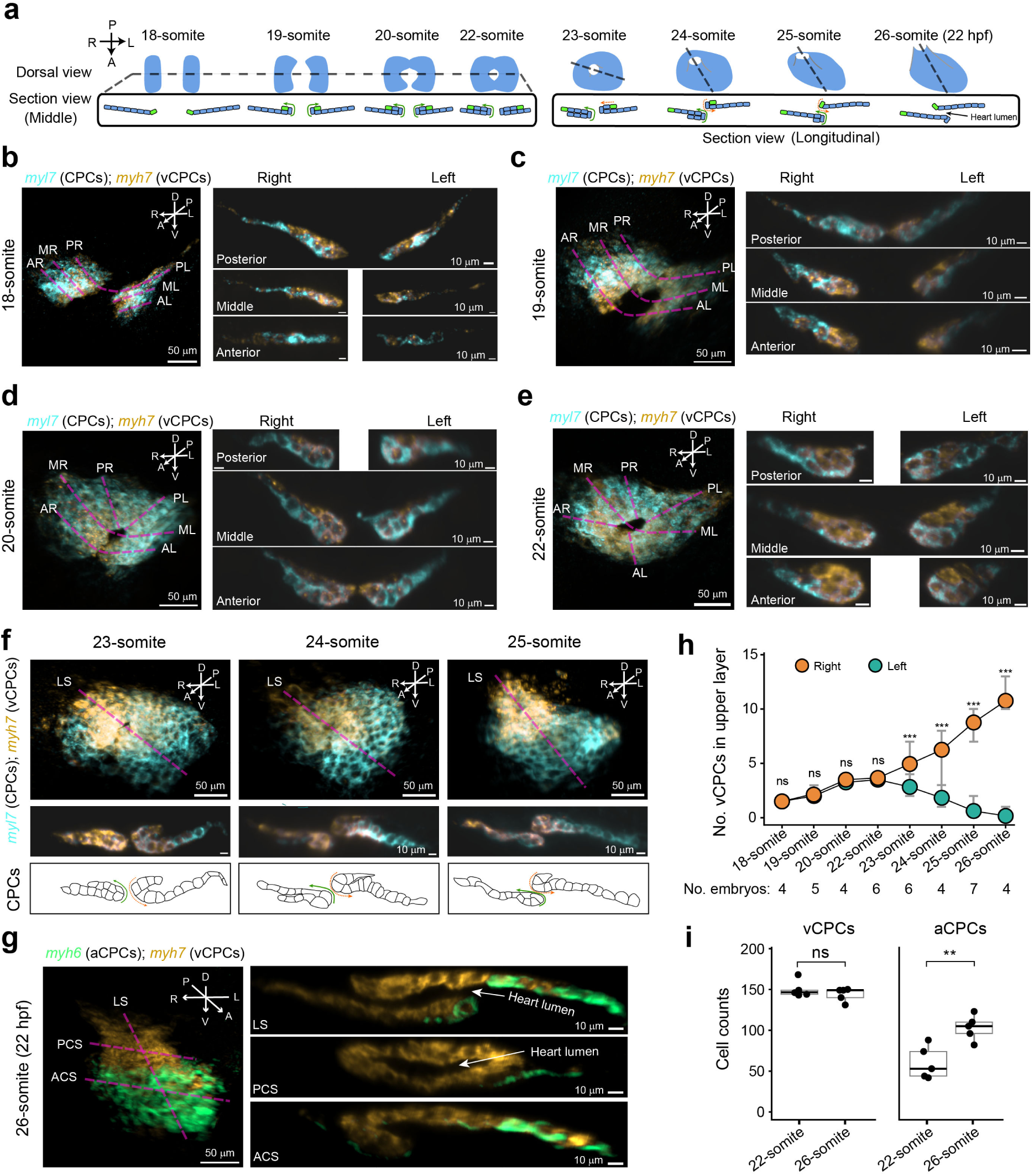
Characterization of ventricular cardiomyocyte precursor involution during their medial migration and extension process. **a**, Scheme of acquisition of sections shown in **b-g**. Directions: A, anterior; P, posterior; L, left; R, right; D, dorsal; V, ventral. The medial most vCPCs were in green. Green arrows indicated involution direction and dashed orange arrows indicated retraction direction. By 22 hpf, CPC sheets becoming single layer again by retraction and involution movement, forming heart lumen. **b-g**, Images of the cardiac cone and transverse optical sections at indicated stages. FISH were performed for the general cardiomyocyte precursor (CPC) marker *myl7,* the vCPCs marker *myh7*, and aVCPCs *myh6* as indicated. The whole cardiac cone was shown on the left (**b-e, g**) or top (**f**) and the optical sections at indicated positions were shown on the right (**b-e, g**) or bottom (**f**) panels. In (**g**), CPCs were outlined with sheet movement directions indicated (as defined in (**a**)) in the bottom panel. AR, anterior right; MR, middle right, PR; posterior right; AL, anterior left; ML, middle left, PL; posterior left; LS, longitudinal section; ACS, anterior cross section; PCS, posterior cross section. **h,** The number of vCPCs in the upper layer on different sides and at different stages. Circular dots denoted the mean values, while error bars showed the maximum and minimum values. Three optical sections per embryo were counted. Difference between left and right number at each stage was analyzed by paired two-tailed Student’s *t*-test. ns, nonsignificant, *P* > 0.05; * *P* < 0.05; ** *P* < 0.01; *** *P* < 0.001. **i**, The number of vCPCs and aCPCs in the whole cardiac cone at indicated stages. Embryos were labeled by FISH with *myh6* and *myh7* probes plus DAPI staining. The *myh6*- or *myh7*-positive cells were counted by 3D light-sheet microscopic observations. Each dot represented cell counts of one embryo. Difference between the two stages was statistically analyzed by unpaired two-tailed Student’s *t*-test with ns for nonsignificance (*P* > 0.05) and ** (*P* < 0.01).

During subsequent heart tube extension, we saw distinct movement patterns of left and right vCPCs. Right vCPCs continued to involute from 23-somite to 26-somite stage, causing bottom-layer CPCs to gradually move to the upper layer (Fig. 1f-h). In contrast, left vCPCs ceased involution and initiated retraction, allowing upper-layer cells to return to the lower layer (Fig. 1f-h). Nevertheless, these two different movement patterns eventually resolved the double-layered structure back into a single layer and created the heart lumen with left vCPCs on the dorsal side and right vCPCs on the ventral side (Fig. 1g). Together, these observations suggest that left and right vCPCs undergo different movements during late stages of lumen formation.

Another previously unsolved question is whether CPCs are undergoing mitosis during involution and extension movements. Light-sheet microscopic and confocal microscopic observations following FISH found neither any mitotic vCPCs nor mitotic aCPCs though mitotic non-CPC cells were observed (Extended Data Fig. 2). Based on 3D light-sheet microscopic observations, we found that the number of *myh7*-positive vCPCs in the whole cardiac cone remained comparable between 22-somite and 26-somite stages (Fig. 1i, left panel). However, the number of *myh6*-positive aCPCs increased at the 26-somite stage compared to that at the 22-somite stage (Fig. 1i, right panel), which might be due to the emergence of newly specified aCPCs with initiation of *myh6* expression. These observations suggest that vCPCs do not proliferate during cardiac cone formation.

### vCPCs undergo a dynamic reorientation of polarity during involution

Our next question was whether dramatic cell rearrangements during vCPC involution were accompanied by cell polarity remodeling. To address this issue, we performed immunofluorescence (IF) for a subset of tight junctional and adhesion proteins using dissected trunk parts of 16- to 22-somite stage *Tg(myl7:EGFP)* transgenic embryos, in which the cardiomyocytes were labeled by EGFP^40^, followed by confocal microscopy (Extended Data Fig. 1b). The results revealed that ZO-1 (also named TJP1), a tight junction adaptor protein, was clearly localized at both inner (apical) and outer (basal) sites of junctions between adjacent vCPCs in the single-layer sheet (Fig. 2a), a pattern distinct from the classical apical-basolateral polarity typical of epithelial cells. Notably, the signal of ZO-1 at the inner site weakened progressively with vCPCs involution, showing an inverse correlation with the depth of the process. Eventually, ZO-1 localized exclusively to outer junctions, with no detectable signal at the inner surface or the newly formed contact surface (inner interface) between the two vCPC layers. This dynamic redistribution of ZO-1 indicated polarity reorientation of involuting and involuted vCPCs, which stands in stark contrast to the persistent apical-basal polarity of classical epithelial cells.

**Fig. 2.**
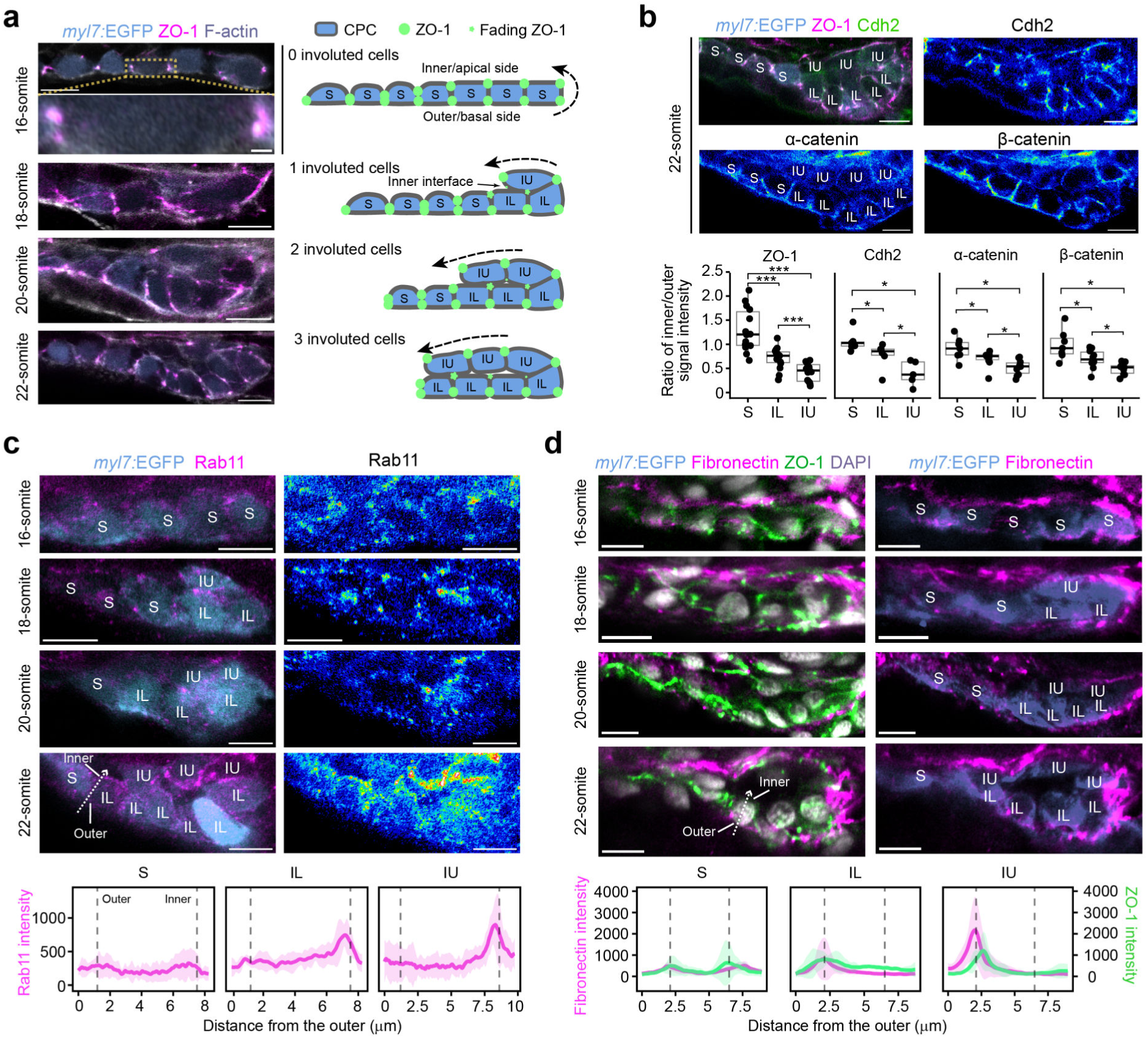
Dynamic location of junctional and adherens proteins in vCPCs during involution. *Tg(myl7:EGFP)* transgenic embryos, in which EGFP were expressed in all cardiomyocytes, were used for immunofluorescence, followed by confocal microscopy (see procedure in Extended Data Fig. 1). One high-quality optical transverse section from a single embryo, with the left or right CPC sheet located roughly at the middle of the cardiac cone, was chosen as the representative image (same selection for subsequent figures). **a,** ZO-1 distribution. F-actin was co-stained to outline cell borders. Left, representative ZO-1 IF images; right, illustration of CPC involution and ZO-1 dynamics. Cell type definition: S, single-layer pre-involution CPC; IL, involuting CPC in the lower layer (IL); IU, involuted CPC in the upper layer. **b**, Distributions of ZO-1, Cdh2, α-catenin and β-catenin. For better view, signals of Cdh2, α-catenin and β- catenin were shown in heatmap. Lower panel, ratio of inner/outer signal intensity in different cell types. Each dot represented an individual cell. Data were from >5 independent embryos. Unpaired two-tailed Student’s *t*-test was performed. *, *P* < 0.05; ***, *P* < 0.001. **c**, **d**, Distributions of Rab11 (**c**) and Fibronectin (**d**). The top right panel in (**c**) showed heatmap of Rab11 signals. Bottom panels, distribution of indicated signals in different cell types across the outer-inner axis (indicated by dashed arrows in the top panel) at 16- to 22-somite stage. Lines and shaded areas represented the mean intensity± s.d.. Five cells from different embryos were measured.

Reorientation of cell polarity is a fundamental cellular mechanism underlying tissue and organ morphogenesis^41–43^. To quantify vCPCs polarity reorientation clearly, we classified vCPCs into three groups: single-layer pre-involution cells (S), involuting cells in the lower layer (IL), and involuted cells in the upper layer (IU) (Fig. 2a, right panel) and examined other adhesive junction components. The results showed that N-Cadherin (Cdh2), α-catenin, and β-catenin signals progressively weakened at inner junctions while remaining consistently high at outer junctions in involuted vCPCs as involution proceeded (Fig. 2b, upper panel), which resembled the dynamic of ZO-1 (Fig. 2a). Statistical analysis of the inner/outer junctional intensity ratio for S, IL, and IU cell groups confirmed the decrease of those signals at the inner junctions (Fig. 2b, lower panel). These observations indicate that vCPCs reorient their polarity and adhesion during the involution process.

To look for upstream candidates regulating vCPC polarity establishment and maintenance, we first examined dynamic location of Rab11, an important regulator of targeted delivery of cell adhesion proteins ^44,45^. Immunofluorescence revealed no asymmetrical enrichment of Rab11 in pre-involution vCPCs but obvious enrichment beneath the plasma membrane on the inner side in IL and IU vCPCs (Fig. 2c), suggesting its potential implication in regulation of vCPCs polarity remodeling during involution. Then, we examined distribution of Fibronectin, a component of the extracellular matrix (ECM), which is known to play a critical role in maintaining ZO-1 localization^28^. The result showed that Fibronectin was enriched in the inner-side ECM of single-layered vCPCs and progressively increased in the outer-side ECM of IL and IU vCPCs during involution, whereas it was absent at the interface between the upper and lower vCPC layers (Fig. 2d). Notably, the dynamics of Fibronectin deposition in vCPCs closely coincided with changes in ZO-1 levels. Thus, it is likely that Fibronectin-marked ECM may play a role in remodeling of vCPCs polarity during involution.

### Podxl drives de-adhension at the inner surface of involuting and involuted vCPCs

Our next question was how junctional and adhesive molecules are cleared from the inner surface of involuting and involuted vCPCs. Podocalyxin (Podxl) is a highly sialylated apical membrane protein that drives removal or relocation of junctional complexes during morphogenetic processes^46–48^. A previous study reported that *podxl* knockdown in zebrafish embryos resulted in pericardial edema with polarity defects^33^, suggesting its involvement in cardiogenesis. So then, we explored the potential role of Podxl in de-adhesion of involuting and involuted vCPCs. Due to lack of suitable Podxl antibody, we injected *podxl-mCherry* mRNA into one-cell stage embryos and observed localization of Podxl-mCherry in vCPCs during involution by immunofluorescence of mCherry. We observed that, prior to involution, Podxl-mCherry was predominantly localized to intracellular vesicles, with no evident enrichment at the membrane or polarized distribution (Fig. 3a, top panel). Following involution, Podxl-mCherry became obviously redistributed to the inner membrane surface, accumulating specifically at the contact interface between upper and lower vCPCs (Fig. 3a, middle and bottom panels). When *podxl* was knocked down using an antisense morpholino (MO)^33^, as expected, clearance of ZO-1 and Cdh2 from the inner surface between IL and IU vCPCs was impaired, resulting in their aberrant retention at the interface (Fig. 3b). This result indicated an essential role of Podxl in removing junctional and adherens proteins in vCPCs.

**Fig. 3.**
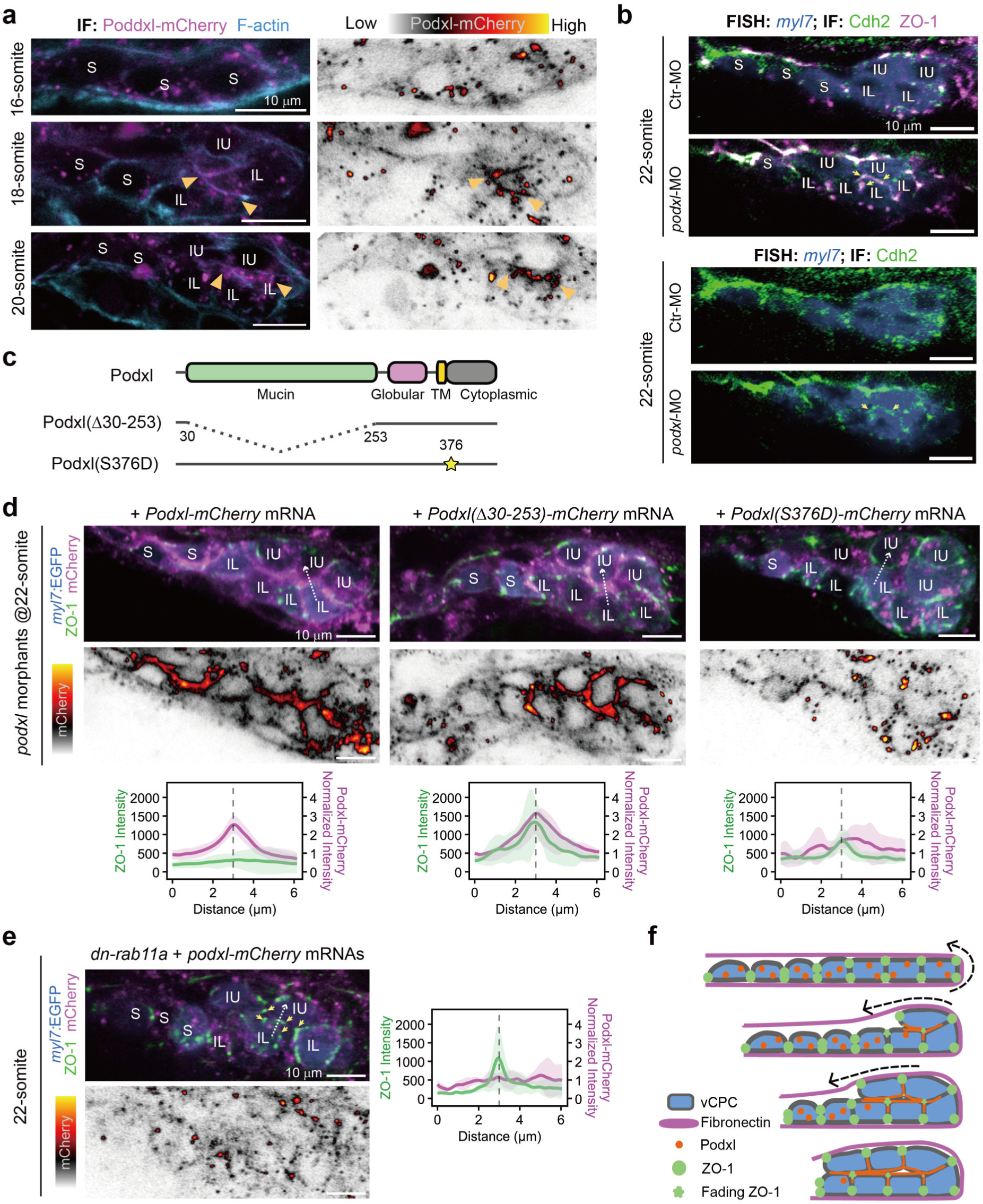
Podxl translocates to the inner surface to clear adhesions during polarity reorientation. **a,** Enrichment of Podxl-mCherry at the inner suface. *podxl-mCherry* mRNA was injected into one-cell stage *Tg(myl7:EGFP)* embryos at 100 pg per embryo, following by IF and confocal microscopy at indicated stages (see procedure in Fig. S1). Single optical section was shown. The heatmaps of Podxl-mCherry signals were shown in the right panel. The inner interface was indicated by orange arrowhead. **b**, Retention of ZO-1 and Cdh2 at the inner interface of vCPCs in *podxl* morphants. One-cell stage WT embryos were injected with 0.2 nM of *Podxl*-MO or control MO (ctr-MO) per embryo and subjected to IF and FISH at the 22-somite stage. The retained ZO-1 (top panel) and Cdh2 (bottom panel) signals at the inner interface were indicated by arrows. **c,** Illustration of domain composition of WT and mutant Podxl. The positions of critical residues were numbered. **d**, Alteration of ZO-1 localization in vCPCs of embryos injected with different podxl mRNAs. One-cell stage WT embryos were injected with indicated mRNA (100 pg/embryo) and subjected to IF for ZO-1 and mCherry. Top panel, IF images; middle panel, heatmap of mCherry signals; bottom panel, intensity distribution of indicated signals across the inner interface (indicated by dashed arrows in top panel). The intensity was the mean ± SD from 5 embryos per group. e, effect of Rab11 inhibition on ZO-1 and Podxl localization. One-cell stage WT embryos were injected with *dnrab11*, encoding dominant negative form of Rab11, and *podxl-mCherry* mRNAs, each at 100 pg/embryo and subjected to IF for ZO-1 and mCherry. The arrows in the left top panel indicated ectopic ZO-1 signals at the inner junctions. Left lower panel, heatmap of Podxl-mCherry signal; right panel, intensity distribution of indicated signals across the inner interface (indicated by dashed arrows in left top panel). The intensity was the mean ± s.d. from 5 embryos. **f**, Illustration of ZO-1 and Podxl in vCPCs during involution.

To identify functional domains of Podxl, we performed rescue experiments by coinjecting *podxl*-MO with different isoforms of Podxl fused to mCherry (Fig. 3c). Coinjection of *podxl-mCherry* mRNA with *podxl*-MO re-eradicated ZO-1 accumulation at the interface in *podxl* morphants, suggesting a rescue effect (Fig. 3d, left panel). In contrast, overexpression of *podxl(Δ30-253)-mCherry* mRNA, which encodes Podxl mutant isoform lacking the mucin domain for anti-adhesive activity^49,50^, failed to prevent ZO-1 accumulation at the interface in *podxl* morphants although this mutant protein was correctly translocated to the inner membrane surface of involuted vCPCs (Fig. 3d, middle panel). Another Podxl mutant isoform, Podxl(S376D)-mCherry, which was a phosphomimetic equivalent of rat Podxl (S415D) mutant blocking RhoA and Ezrin activation^47,51^, was stabilized in intracellular vesicles of involuting vCPCs upon overexpression in *podxl* morphants and unable to displace ZO-1 at the interface (Fig. 3d, right panel). Furthermore, expression of a dominant-negative Rab11a (S25N)^52,53^ in WT embryos, which was expected to disrupt Podxl polarized trafficking, impaired Podxl-mCherry enrichment at the inner interface and resulted in retention of inner junctional ZO-1 (Fig. 3e), mimicking the effect of the phosphomimetic Podxl(S376D) overexpression. These results suggest that Podxl is necessary for de-adhesion of vCPCs between the double layers to facilitate movements of involuted vCPCs (Fig. 3f), including subsequent retraction and extension.

### Deficiency of Afdna leads to dysfunctional heart with circulatory stasis

Afadin (AFDN/AF6), a central apical junction scaffold protein, coordinately controls cell–cell adhesion by regulating adherens junctions via nectin binding and linkage to the F-actin cytoskeleton and governs epithelial polarity by recruiting core polarity complexes downstream of Rap1 and Ras GTPases^48,54,55^. However, its implication in heart morphogenesis has been unexplored. We cloned *afdna,* one of the zebrafish orthologs of mammalian *Afdn.* Whole-mount in situ hybridization (WISH) revealed that *afdna* transcripts are present ubiquitously in early embryos from the 1-cell stage to 24 hpf (Extended Data Fig. 3a-f) and apparently in the ventricle of the heart at 48 hpf (Extended Data Fig. 3f-g). To examine subcellular localization of endogenous Afdna protein, we generated a custom antibody against zebrafish Afdna. Immunofluorescence using this antibody detected apical enrichment of Afdna in neuroepithelia at the 16-somite stage, otic vesicle epithelia at 24 hpf, and optic vesicle epithelia at 24 hpf (Extended Data Fig. 4a-c), which resemble localization of mouse Afadin in these cell types^56^. In contrast, Afdna was largely colocalized with ZO-1 in vCPCs, including at the inner/apical and outer/basal junctions of single-layer pre-involution vCPCs (16-somite stage), the outer and lateral membrane of double-layer vCPCs (22-somite stage), and also outer and lateral membrane of vCPCs within the lumenized ventricle (24 hpf) (Fig. 4a). Thus, we hypothesized that Afdna might play a critical role in zebrafish ventricle morphogenesis through distinct mechanisms.

**Fig. 4.**
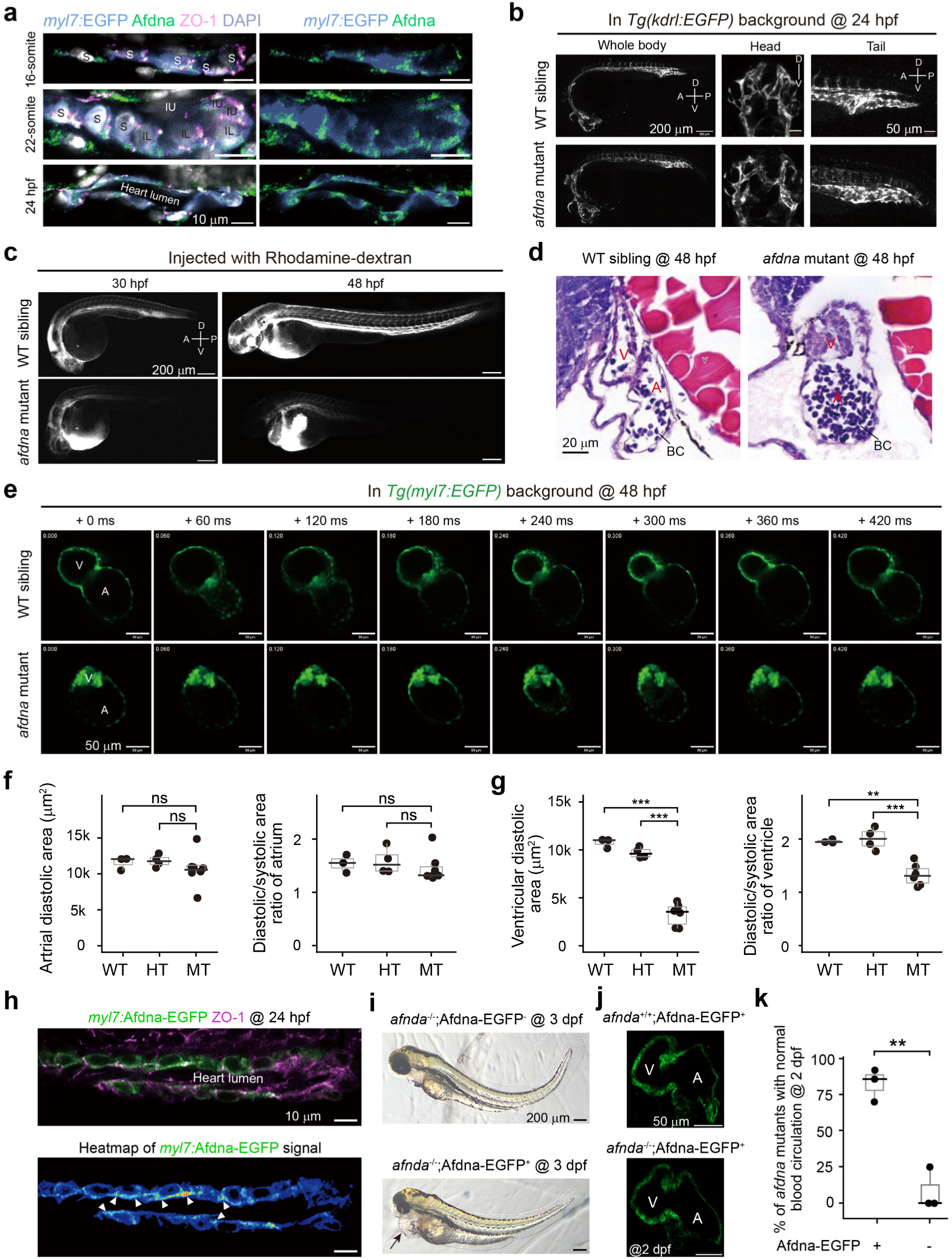
Afdna is required for ventricular development. **a,** Enrichment of Afdna in outer surface of involuted vCPCs. Representative confocal images were shown with cell types indicated. **b**, Visualization of vascular network. Embryos were derived from crosses between *afdna* heterozygotes and *Tg(kdrl:EGFP)* transgenic fish with vasculature labeled. Embryo orientation: A, anterior; P, posterior, D, dorsal; V, ventral. **c**, Examination of circulation. Rhodamine-dextran beads was injected into the common cardinal vein (CCV) of embryos and observed by confocal microcopy. **d**, Anatomical structure of the heart. Embryos were sectioned and stained with Hematoxylin and Eosin. A, atrium; V, ventricle; BC, blood cells. **e**, Time-lapse montages of heart morphology. Embryos were derived from crosses between *afdna* heterozygotes and *Tg(myl7:EGFP)* transgenic fish in cardiomyocytes labeled, observed by confocal microscopy. The remanent embryonic tissue after microscopy was used for genotyping (same for other figures). Single optical sections were shown. A, atrium; V, ventricle. **f, g,** Diastolic area and ratio of diastolic to systolic area of atrium (**f**) and ventricle (**g**). Each dot represented one embryo. Unpaired two-tailed Student’s *t*-test was performed. ns, nonsignificant; **, *P* < 0.01; ***, *P* < 0.001 (same applied to (**j**)). **h**, Afdna-EGFP enrichment in inner surface and lateral junctions (indicted by arrowheads) of cardiomyocytes in *Tg(myl7:afdna-EGFP)* transgenic embryos. **i**, A representative *afdna* mutant embryo with Adfna-EGFP expression, which was derived from an incross of *afdna;Tg(myl7:afdna-EGFP)* heterozygous fish. Note the absence of pericardia edema (indicated by an arrow) in a rescued embryo. **j**, Restoration of ventricular chamber in a representative mutant with transgenic Afdna-EGFP expression. **k**, Ratio of *afdna* mutants with normal blood circulation in *Tg(myl7:afdna-EGFP)* transgenic background. Each dot represented one batch (≥ 40 embryos per batch). ** *P* < 0.01.

To explore Afdna function, we generated the *afdna* mutant line *afdna^tsu-^*^5^ using CRISPR/Cas9 technology. The mutant allele carries a 5-bp deletion in the first exon, resulting in a premature stop codon (Extended Data Fig. 5a). The absence of Afdna protein in *afdna* homozygous mutants, as examined by Western blot (Extended Data Fig. 5b), indicated that the mutant allele is a null allele. Mutant embryos developed without obvious morphological defects before blood circulation started (24 hpf) (Extended Data Fig. 5c). By 48 hpf, however, 70-100% of mutants, varying among fish pairs, displayed obvious pericardial edema with heart beating (Extended Data Video 1) and dorsal curvature of the trunk (Extended Data Fig. 5c). Mutants died around 6 days postfertilization (dpf). Examination of vasculature in *afdna* mutants in *Tg(kdrl:EGFP)* background at 24 hpf, in which the blood vessels were labeled by EGFP^57^, revealed no obvious anomalies (Fig. 4b), but injected Dextran rhodamine dye was unable to circulate in the mutants (Fig. 4c). Then, we shifted to carefully examine the heart structure in mutants. Hematoxylin and Eosin staining showed that, in mutants, the atrial chamber looked normal but with stacked blood cells, whereas the ventricle region failed to establish a proper chamber architecture and displayed multilayered cardiomyocytes (Fig. 4d). Live imaging of *afdna;Tg(myl7:EGFP)* embryos at 48 hpf by confocal microscopy also showed that the atrial chamber of mutants had normal morphology with normal diastole and systole (Fig. 4e, f), whereas the anterior mutant ventricle chamber had a small canal flanked by cardiomyocyte masses and beat laboriously (Fig. 4e, g, and Extended Data Video 2). Taken together, these observations support the idea that *afdna* is necessary for the ventricle chamber formation.

To test cell-autonomous function of *afdna* in heart development, we made the *Tg(myl7:afdna-EGFP)* transgenic line, which expressed Afdna-EGFP fusion protein specifically in cardiomyocytes (Fig. 4h) and did not cause detectable defects in WT genetic background. Then, *Tg(myl7:afdna-EGFP)* fish were crossed to *afdna* heterozygotes to generate *afdna^+/-^;Tg(myl7:afdna-EGFP)* fish and subsequent *afdna^−/-^;Tg(myl7:afdna-EGFP)* transgenic mutant embryos. We found that *afdna* mutants with transgenic expression of Afdna-EGFP could reduce the pericardial edema (Fig. 4i) and recover ventricular chamber (Fig. 4j). Importantly, transgenic expression of Afdna-EGFP successfully restored blood circulation in the majority of mutants (Fig. 4k). Therefore, Afdna regulates the heart ventricle formation in a cell-autonomous fashion.

### Afdna deficiency impairs polarity reorientation during vCPCs involution and blocks ventricular region extension

To investigate developmental origin of ventricular chamber malformation in *afdna* mutants, we examined expression patterns of heart lineage markers at various stages by FISH. The expression pattern of the cardiogenic mesoderm marker *nkx2.5* at the 14-somite stage and the CPC marker *myl7* at the 18-somite stage was unaltered in *afdna* mutants (Fig. 5a), suggesting a correct specification of CPCs. Examination of the vCPCs marker *myh7* and the aCPCs marker *myh6* at the 16- and 20-somite stages indicated that both vCPCs and aCPCs in *afdna* mutants could initiate medial migration and form the cardiac cone (Fig. 5b). After the 21-somite stage, both aCPC sheet and vCPC sheet of the cardiac cone in wild-type (WT) embryos initiated extension to form a linear canalized heart tube. In contrast, the vCPC sheet in mutants appeared unable to properly extend, and as a result, formed the ventricle with an abnormally narrow chamber, although the aCPC sheet extended normally to form the atrium with a normal chamber (Fig. 5b). Besides, the leftward tilting of the cardiac cone in *afdna* mutants was observed, although completion of this process was slightly delayed compared to WT embryos (Fig. 5b). These observations confirm the specific ventricular chamber defect in *afdna* mutants and imply that this defect may associate with improper involution and impaired retraction and extension of involuted vCPCs in mutants.

**Fig. 5.**
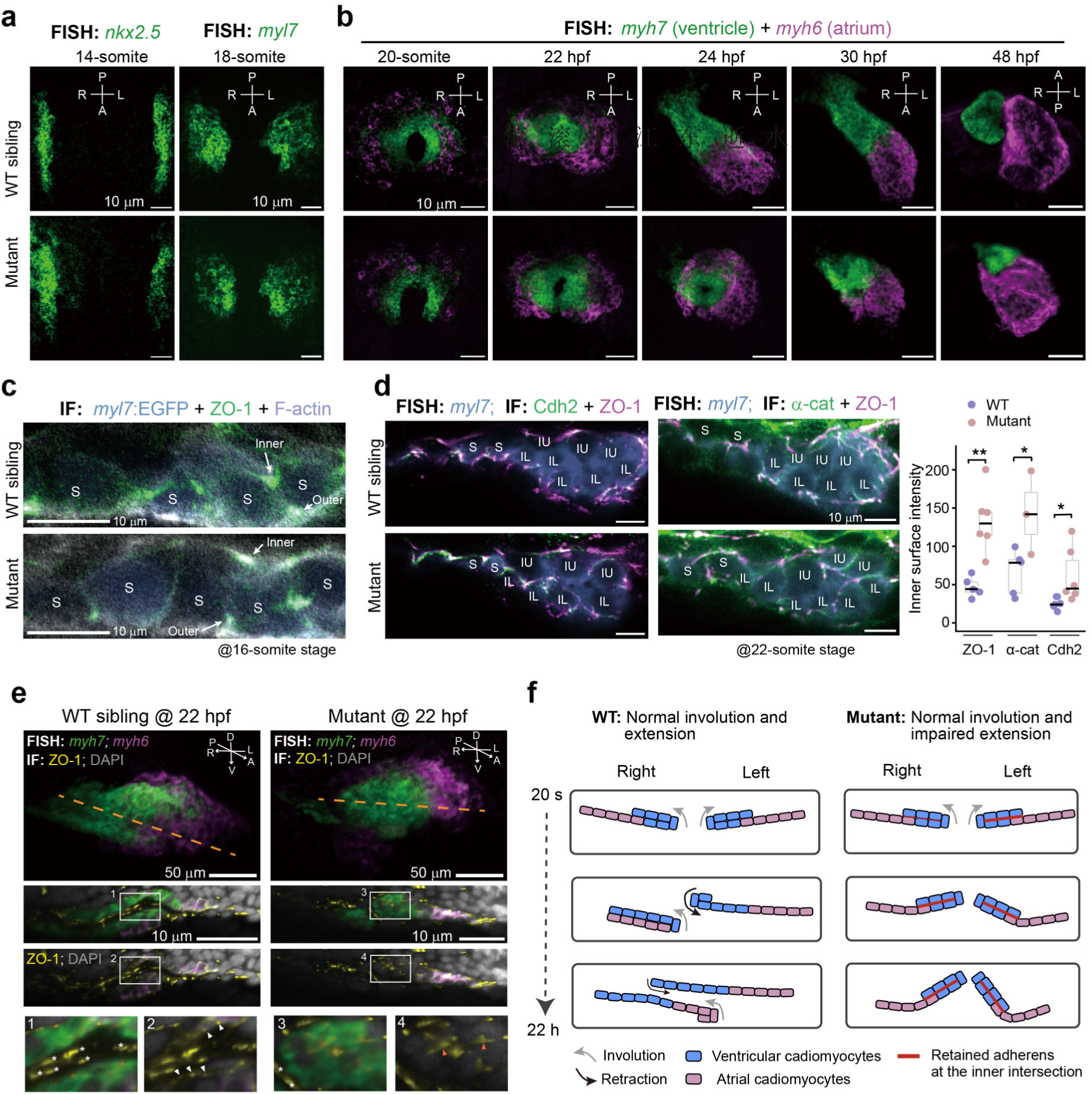
Defective polarity and involution/extension of the cardia cone in *afdna* mutants. **a**, FISH detection of the cardiac mesoderm marker *nkx2.5* (left) and the cardiomyocyte marker *myl7* (right) in *afdna* mutants and WT siblings. Embryos were dorsally viewed by confocal microscopy. A, anterior; P, posterior; L, left; R, right. **b**, Time-lapse montages of cardia cone/heart tube extension in *afdna* mutants and WT siblings with *Tg(myl7:EGFP)* transgenic background. FISH was performed for the ventricle marker *myh7* and the atrial marker *myh6* at indicated stages. Z-stack confocal images were shown. Note that the ventricular chamber of *afdna* mutants was abnormal. **c**, Normal localization of ZO-1 in single-layer non-involuting vCPCs at the 16-somite stage. Embryos with *Tg(myl7:EGFP)* transgenic background were subjected to IF. **d**, Abnormal localization of Cdh2, ZO-1 and α-catenin (α-cat) at the interface between IU and IL vCPCs. Embryos were subjected to FISH for *myl7* and IF for ZO-1/Cdh2 (left) or ZO-1/α-catenin (right), followed by confocal microscopy. Right graph, quantification of adhesion components at inner contact interface; each dot represented an independent embryo; statistical significance of the difference was determined by unpaired two-tailed Student’s *t*-test with \**P* < 0.05 and ** *P* < 0.01. **e**, Abnormal ZO-1 localization and defective retraction of involuted vCPCs in *afdna* mutants. Embryos were simultaneously subjected to FISH for *myh7* and *myh6* and IF for ZO-1, followed by light-sheet microscopy. Top panel, whole image of the cardiac cone; second panel, optical section view along the line indicated in the top panel; third panel, same as second pane except removed *myh7* signal; bottom panel, enlarged areas boxed in the second and third panels. ZO-1 signals at the outer junctions of single-layer vCPCs in WT embryo and at the inner interface in *afdna* mutant were indicated by white and orange arrowheads, respectively. ZO-1 signals in endocardial cells were indicated by *. F, model showing defective retraction and extension of involuted vCPCs. The gay arrow and black arrow indicated retraction and extension direction, respectively.

Then, we investigated dynamics of vCPCs polarity during involution in mutants. Immunofluorescence revealed normal localization of ZO-1 in inner and outer junction sites of pre-involution single-layer vCPCs in *afdna* mutants at the 16-somite stage (Fig. 5c). By the 22-somite stage, however, tight junctional and adhesion molecules, including ZO-1, Cdh2, and α-catenin, had still accumulated at the inner surface of involuted vCPCs in mutants, which were in contrast to their absence in WT embryos (Fig. 5d, e). Light-sheet microscopy revealed that, in WT embryos at 22 hpf, vCPCs became a single layer again after extension with ZO-1 exclusively localized to the outer junctions (Fig. 5e, f, left panels), whereas vCPCs in mutants still had double layers with preserved ZO-1 at the inner junctions (Fig. 5e, f, right panels). The defects in clearance of Cdh2 and ZO1 at the inner interface between upper and lower vCPCs layers in mutants were also confirmed by confocal microscopy at 22 and 24 hpf (Extended Data Fig. 6a). Abnormal ZO-1 localization in disorganized cardiomyocyte mass was still obvious in the ventricle of mutants at 48 hpf (Extended Data Fig. 6b). These observations support the idea that Afdna participates in clearance of adhesions between two layers of vCPCs, facilitating vCPC movements during heart cone elongation and ventricular chamber formation.

### Afdna retains activated Rap1 and Cdc42 at the outer adherens

To further delineate the molecular mechanism underlying polarity reorientation, we examined distribution of several key regulators in *afdna* mutants. First, immunofluorescence imaging revealed that the distribution patterns of Fibronectin and Rab11 in vCPCs of *afdna* mutants during involution resembled those in WT embryos (Extended Data Fig. 7), signifying that Afdna may not regulate vCPC polarity remodeling via ECM or Rab11. Compared to WT embryos, however, *afdna* mutants showed less prominent enrichment of ectopically expressed Podxl-mCherry in the inner plasma membrane of involuted vCPCs (Fig. 6a), which suggests that Afdna is required for Podxl translocation to the inner-side cytoplasm of involuted vCPCs.

**Fig. 6.**
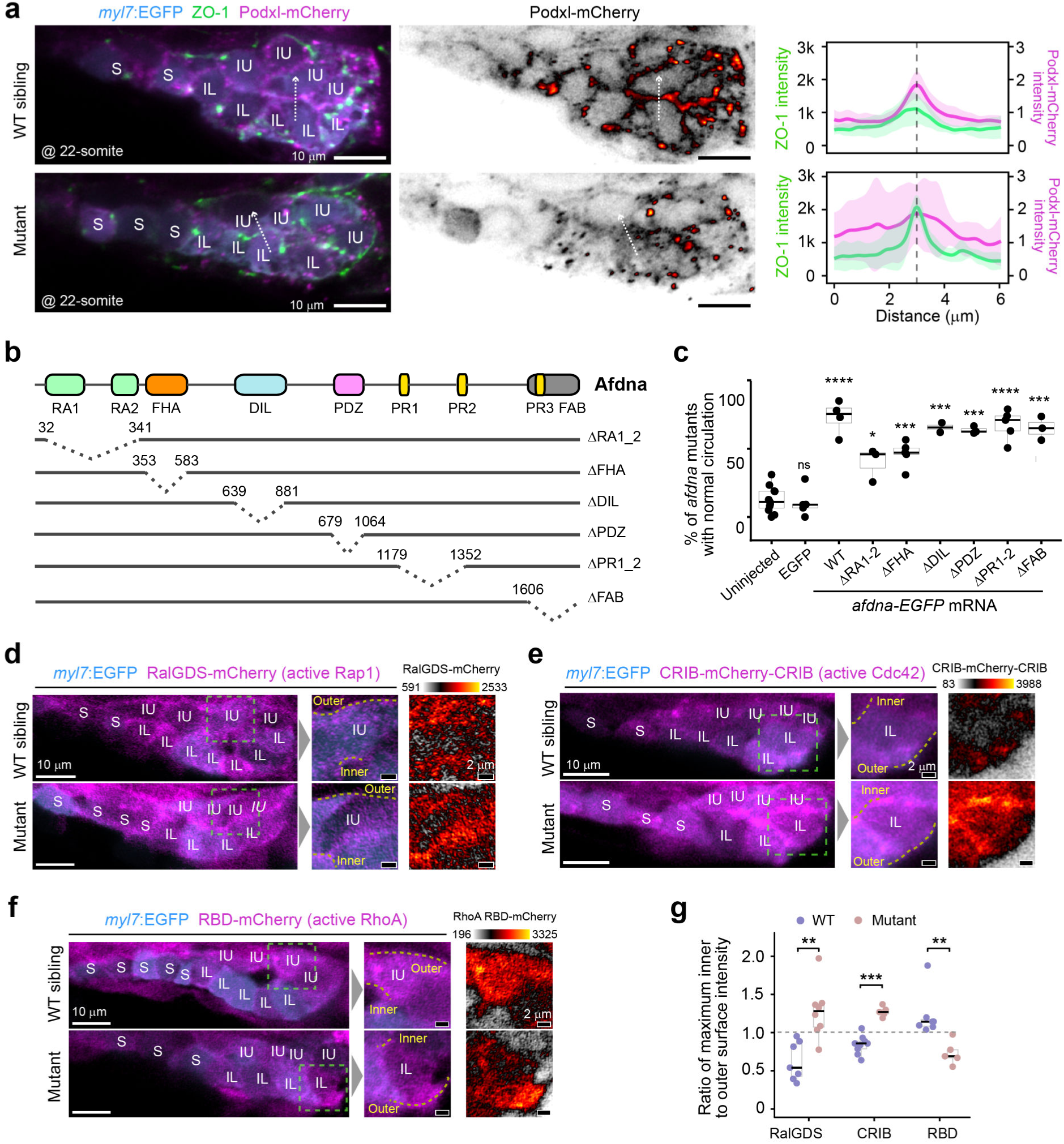
Afdna retains activated Rap1 at the outer adherens. **a,** Defective targeting of Podxl to the inner interface of vCPCs in *afdna* mutants. Embryos in *Tg(myl7:EGFP)* transgenic background were injected with *podxl-mCherry* mRNA (100 pg per embryo) at the one-cell stage and immunostained for ZO-1, mCherry, and EGFP, followed by confocal microscopy. Left, single optical sections with all examined signals; middle, heatmap of mCherry; right, Podxl-mCherry and ZO-1 fluorescence intensity across the inner contact surface as indicated by dashed arrows in the left panel. Data were mean intensity ± SD from 5 embryos. **b**, Illustration of Afdna protein domain composition of WT Afdna and domain-deletional variants. **c**, Rescue effects of different forms of Afdna variants fused to EGFP on blood circulation in *afdna* mutants. Embryos from *afdna* heterozygote incrosses were injected at the 1-cell stage with indicated mRNA species (400 pg per embryo) and observed for blood circulation at 2 dpf, followed by genotyping to identify *afdna* mutants. Each dot represented one batch of embryos. Data were mean ± s.d.. Injected groups each was compared to uninjected group. Unpaired two-tailed Student’s *t*-test. ns, nonsignificant, *P* > 0.05; * *P* < 0.05; *** *P* < 0.001; ****, P < 0.0001. **d-f**, Altered localization of active Rap1 (**d**), Cdc42 (**e**) and RhoA (**f**). Embryos from *afdna* heterozygote (with *Tg(myl7:EGFP)* background) incresses were injected with 50 pg (per embryos) mRNA encoding an indicated GTPase sensor at the one-cell stage and immunostained for mCherry and EGFP at the 22-somite stage, followed by confocal microscopy. An area over the inner interface between IU and IL vCPCs was enlarged. **g**, Altered ratio of inner to outer signal intensity for each activated GTPase. Each dot represented the data from one embryo and at least 5 embryos for each group were analyzed. Statistical significance of the difference was determined by unpaired two-tailed Student’s *t*-test. \*\**P* < 0.01; *** *P* < 0.001.

Next, we adopted a domain-mapping approach to define functional motifs of Afdna in vCPC polarity remodeling during involution (Fig. 6b). mRNAs encoding deletional variants fused to EGFP were individually injected into one-cell stage embryos from *afdna* heterozygous intercrosses. After genotyping, the proportion of mutant embryos with blood circulation at 2 dpf was calculated. We observed that, while overexpression of any mutant variants was able to restore blood circulation in mutant embryos, the efficacy was consistently lower than that observed following overexpression of the full-length *afdna-EGFP* (Fig. 6c). Of note, overexpression of mutant Afdna with deletion of Ras-associated domains 1–2 (RA1–2) or the forkhead-associated domain (FHA) showed a significantly reduced rescue efficacy, indicating that the RA and FHA domains are important for Afdna regulation of heart development.

The RA1–2 domain of Afadin is known to bind Ras and Rap GTPases^58–60^. Rho GTPases have been demonstrated to antagonize Rap GTPases reciprocally in adhesion regulation^61,62^. Then, we asked if loss of Afdna in vCPCs might be linked to dysregulation of Rap GTPases and Rho family GTPases. Owing to impractical biochemical analysis for low number of vCPCs and challenges in live imaging on transverse sections, we used translocation-based probes/sensors to visualize the intensity and localization of active GTPases, including the active Rap1 probe RalGDS-mCherry^63^, the active RhoA probe RhoA RBD-mCherry^64,65^, and the active Cdc42 probe mCherry-CRIB-mCherry^66^. Synthesized mRNAs encoding these probes were individually injected into one-cell stage embryos and the embryos at the 22-somite stage were then imaged by confocal microscopy to quantify the mCherry intensity ratio between the outer and inner membranes of IU and IL vCPCs. In WT embryos, activated Rap1 and Cdc42 were enriched at the outer surface (Fig. 6d, e, g), whereas activated RhoA level was slightly higher at the inner surface (Fig. 6f, g). In contrast, *afdna* mutants exhibited significantly higher levels of activated Rap1 and Cdc42 at the inner surface of both IL and IU vCPCs, while conversely showing reduced levels of activated RhoA at this same domain. These results suggest that Afdna acts to maintain active Rap1/Cdc42 at the outer adherens, which permits RhoA activation at the inner-side cytoplasm.

We have shown in the above that Podxl enrichment at the inner surface of IU and IL vCPCs is necessary for removing local adherens (Fig. 3). Therefore, we hypothesized that Afdna might promote translocation of Podxl to the inner surface as a spatial regulator of GTPases. To test this idea, we treated *afdna* mutant embryos with the Rap1 inhibitor GGTI-298, the Rho activator Narciclasine, or both, and observed restoration of blood circulation. The results showed that either the Rap1 inhibitor or the Rho activator partially rescued blood circulation of *afdna* mutants and their co-administration gave a better rescue efficacy (Fig. 7a). Furthermore, simultaneous knockdown of *rap1a* and *rap1b* in *afdna* mutants using specific MOs^34^, each at 0.25 nM, restored blood circulation of *afdna* mutants (Fig. 7c), which associated with reappearance of Podxl enrichment at the inner surface (Fig. 7c). Conversely, expression of a constitutively active Rap1a^67^ (CA-Rap1a) (Fig. 7d) or *rhoa* knockdown in WT embryos (Fig. 7e) prevented Podxl enrichment at the inner surface with the retention of ZO-1. Taken together, these observations suggest that Afdna is required for detaining active Rap1 at the outer surface of involuted vCPCs so that RhoA is activated and facilitates Podxl translocation to the inner surface.

**Fig. 7.**
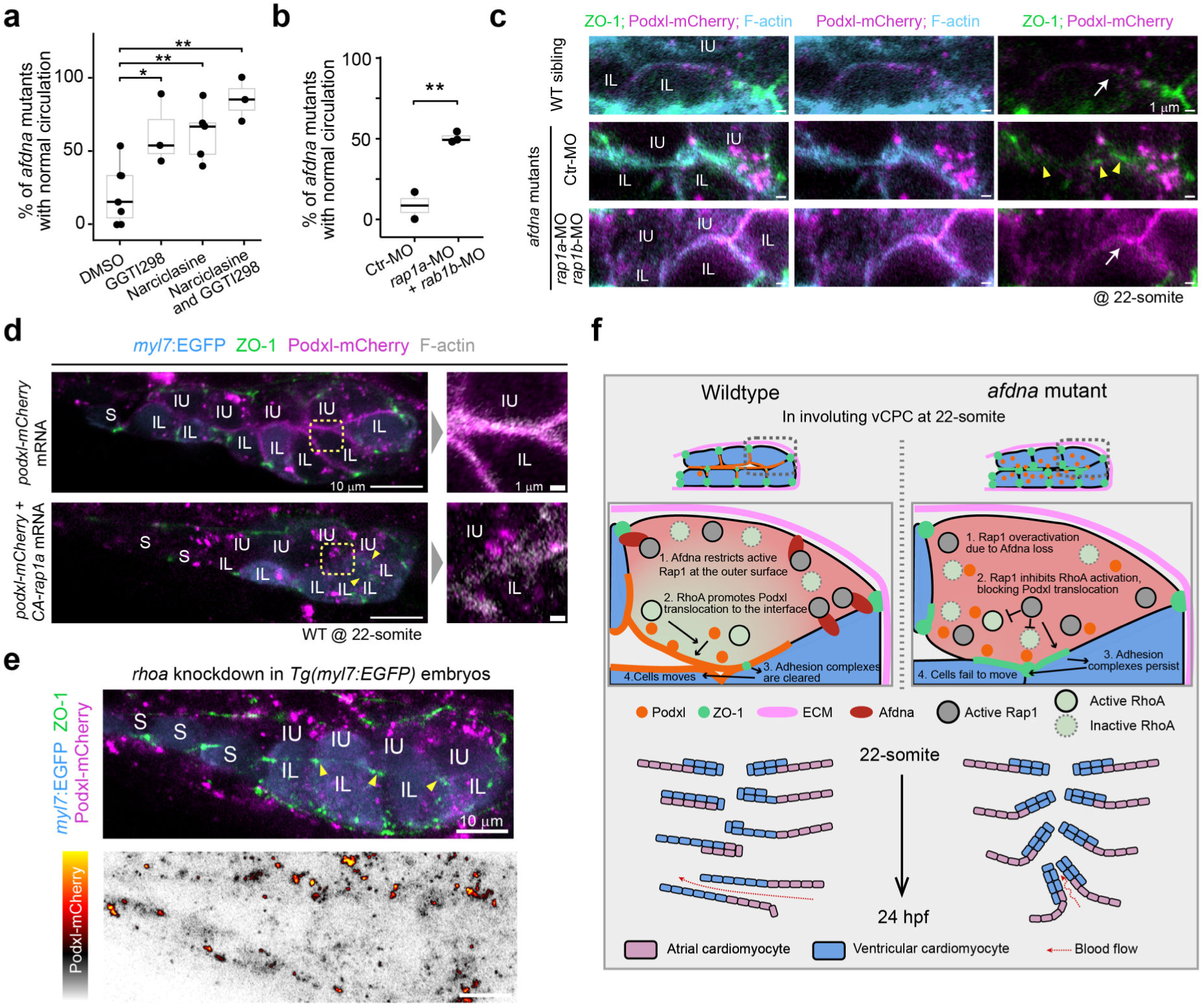
Afdna directs Podxl translocation by regulating distribution of activated GTPases. **a**, Restoration of blood circulation in *afdna* mutants by Rap1 inhibition or/and RhoA activation. Embryos from *afdna* heterozygote incrosses were treated with the Rap1 inhibitor GGTI-298, the Rho activator Narciclasine, or both from 13 hpf to 21 hpf and observed for blood circulation at 2 dpf, followed by genotyping. Each dot represented one batch of *afdna* mutants (n ≥ 20). The data were expressed as mean ± s.d. Unpaired two-tailed Student’s *t*-test was performed with \**P* < 0.05 and ** *P* < 0.01. **b**, co-knockdown of *rap1a* and *rap1b* restores blood circulation of *afdna* mutants. One-cell stage embryos from *afdna* heterozygote incrosses were injected with *rap1a*-MO and *rap1b*-MO or control MO (ctr-MO), each at 0.25 nM, and observed for blood circulation at 2 dpf, followed by genotyping. Each dot represented one batch of mutant embryos (n ≥ 20). The data were expressed as mean ± s.d. Unpaired two-tailed Student’s *t*-test was performed with ** *P* < 0.01. **c**, Corrected distribution of ZO-1 and Podxl in *afdna* mutants with co-knockdown of *rap1a/1b*. Embryos were injected with rap1a-MO/rap1b-MO (as in (b)) and 100 pg podxl-mCherry mRNAs at the one-cell stage and immunostained for ZO-1 and mCherry with phalloidin staining for F-actin at the 22-somite stage. Representative images were single optical sections focusing on involuted vCPCs. Podxl-mCherry enrichment at the inner interface in WT embryos and morphants was indicated by white arrows and ectopic ZO-1 signals at the inner interface were indicated by yellow arrowheads. **d**, Disruption of Podxl location at the inner interface by CA-*rap1a* overexpression. *Tg(myl7:EGFP)* embryos were injected at the one-cell stage with *podxl-mCherry* mRNA alone or together with CA-*rap1a* mRNA (encoding constitutively active Rap1a), each at 100 pg per embryo, and immunostained for EGFP, ZO-1, and mCherry at the 22-somite stage. The boxed areas in the left panel were enlarged in the right panel. The ectopic ZO-1 signals at the inner junctions were indicated by arrowheads (same in (**e**)). **e**, Disruption of Podxl location at the inner interface by *rhoa* inhibition. *Tg(myl7:EGFP)* embryos were injected at the one-cell stage with 100 pg *podxl-mCherry* mRNA and 0.5 nM rhoa-MO and immunostained for mCherry, EGFP, and ZO-1 at the 22-somite stage. The lower image was the heatmap of mCherry intensity, showing the absence of Podxl at the inner interface. **f**, Model of involution and adherens junction clearance of vCPCs in WT and *afdna* mutant embryos (see text for details).

## Discussion

In this study, we redefine the cellular mechanism underlying zebrafish heart tube formation by uncovering a programmed and essential polarity reorientation event during vCPC involution. Contrary to the previous model which implied a static maintenance of polarity, our findings demonstrate that vCPCs undergo a dramatic reprogramming of their polarity axis, which is initiated by asymmetric ECM attachment and executed through the coordinated action of Afdna, small GTPases, and Podxl. We propose a sequential model (Fig. 7f). (1) The asymmetric distribution of ECM, enriched only on the outer surface, provides a directional cue; (2) Afdna, localized at the outer and lateral junctions, functions as a spatial gatekeeper to restrict active Rap1 to the ECM-attached surface; (3) This restriction permits the activation of Rho at the inner, ECM-detached surface; (4) Active Rho promotes the translocation of Podxl to the inner membrane; (5) Podxl clears adhesion complexes (e.g., ZO-1 and Cdh2), enabling the vCPC double layers to slide past one another during subsequent involution (right vCPCs) or retraction/extension (left vCPCs). Disruption of any step in this cascade locks vCPCs in an ectopically adherent state, ultimately preventing ventricular chamber expansion.

During vCPC involution, Fibronectin, one of ECM major components, is removed from the inner surface while increasing at the outer surface in involuting vCPCs (Fig. 2d), suggesting that asymmetric ECM triggers polarity reorientation. However, we have not addressed how this asymmetry arises, which may involve both differential degradation of ECM and polarized secretion. It is known that endothelial cells can secrete Fibronectin asymmetrically and polarize Fibronectin secretion^68^, and that endothelial defects phenocopy heart tube extension failure^69,70^. Further investigation is needed to understand how Fibronectin is polarized, secreted, and degraded.

The ventricle chamber formation undergoes several stages and involves stage-specific polarity reorientation of ventricular cardiomyocytes. Dysregulation of these polarity reorientation events can lead to similar or distinct heart defects. For example, Fibronectin is an ECM trigger of vCPC polarity establishment during medial migration and mediolateral expansion of vCPCs and its mutation in zebrafish *natter* (*nat*) mutants can cause cardia bifida^71^ with disrupted asymmetric localization of the polarity proteins aPKC and ZO-1^28^. Crb2a is a component of the Crumbs polarity complex and its mutation in *oko meduzy* (ome) mutants results in multilayered ventricle wall due to mislocalization of tight and adherens junction proteins during cardiac trabeculation^27^. Similarly, MZ*gpr126^st^*^49^ mutants have multilayered ventricular wall due to the failure of ventricular cardiomyocyte depolarization during trabeculation^72^. In our *afdna* mutants, double-layered vCPCs failed to slide during extension of the cardiac cone due to uncleared tight junctional and adhesion proteins at the interface between two layers, leading to the obliteration of the ventricular lumen with multilayered and disorganized myocardial mass.

The *afdna* mutant phenotype presents the first model of ventricular lumen defect caused by defective vCPC polarity remodeling and extension during the cardiac cone formation. Our study establishes the Afdna-Rap1-RhoA-Podxl regulatory axis of polarity reorientation. It will be interesting to investigate whether this regulatory axis is implicated in the heart morphogenesis of mammals, during which no cardiomyocyte involution happens. Nevertheless, future deep mechanistic studies utilizing *afdna* mutants could dissect the upstream regulatory signals and downstream effector molecules that govern cardiomyocyte stratification and lumen development, ultimately advancing our understanding of the embryonic origins of severe congenital single ventricle defects in human.

Beyond cardiogenesis, our work illustrates a fundamental morphogenetic principle: the dynamic repurposing of cell polarity to drive complex tissue rearrangements. The molecular toolkit we identified, a scaffold protein spatially regulating antagonistic small GTPases to control anti-adhesive factor deployment, may represent a conserved strategy for managing cell cohesion and separation in various developmental contexts. In conclusion, by revising the existing model of heart morphogenesis to include active polarity reorientation, our work not only elucidates a key event in zebrafish development but provides a conceptual framework for understanding the dynamic cellular basis of morphogenesis across evolution.

## Acknowledgements

We thank for the imaging assistance from the Cell Biology Facility of Tsinghua University. We are grateful to other members of the Meng lab for various assistance and insightful discussion.

## Author contributions

Z.W. designed and performed experiments, analyzed data, and write the paper; J.C. helped fish care and breeding; C.P. performed some experiments; X.W. supervised the study; A.M. conceived and supervised the study, analyzed data, and wrote the paper.

